# Embryonic organoids recapitulate early heart organogenesis

**DOI:** 10.1101/802181

**Authors:** Giuliana Rossi, Andrea Boni, Romain Guiet, Mehmet Girgin, Robert G. Kelly, Matthias P. Lutolf

**Affiliations:** Laboratory of Stem Cell Bioengineering, Institute of Bioengineering, School of Life Sciences and School of Engineering, École Polytechnique Fédérale de Lausanne (EPFL), Lausanne, 1015, Vaud, Switzerland; Viventis Microscopy Sàrl, EPFL Innovation Park, building C, Lausanne, 1015, Vaud, Switzerland; Faculté des sciences de la vie, Bioimaging and Optics Platform, École Polytechnique Fédérale de Lausanne (EPFL), Bâtiment AI, Station 15, Lausanne, 1015, Vaud, Switzerland; Aix-Marseille Université, CNRS UMR 7288, IBDM, Marseille, France; Institute of Chemical Sciences and Engineering, School of Basic Science, École Polytechnique Fédérale de Lausanne (EPFL), Lausanne, 1015, Vaud, Switzerland

**Keywords:** Embryonic organoids, cardiogenesis, development, *in vitro* organogenesis, heart, 3D cardiac tissue, cardiac organoid

## Abstract

Organoids are powerful models for studying tissue development, physiology, and disease. However, current culture systems disrupt the inductive tissue-tissue interactions needed for the complex morphogenetic processes of native organogenesis. Here we show that mouse embryonic stem cells (mESCs) can be coaxed to robustly undergo the fundamental steps of early heart organogenesis with an *in vivo*-like spatiotemporal fidelity. These axially patterned embryonic organoids support the generation of cardiovascular progenitors, as well as first and second heart field compartments. The cardiac progenitors self-organize into an anterior domain reminiscent of a cardiac crescent before forming a beating cardiac tissue near a putative primitive gut-like tube, from which it is separated by an endocardial-like layer. These findings unveil the surprising morphogenetic potential of mESCs to execute key aspects of organogenesis through the coordinated development of multiple tissues. This platform could be an excellent tool for studying heart development in unprecedented detail and throughput.

## Introduction

Stem–cell-derived organoids self-organize into complex structures that mimic aspects of the architecture, cellular composition, and function of tissues found in real organs (Clevers, 2016; Lancaster and Knoblich, 2014; Rossi et al., 2018; Sasai, 2013). While most organoids simulate specific features of adult organs, embryonic organoids can capture key processes that occur during early embryonic development, from the pre-implantation blastocyst (Rivron et al., 2018) to early post-implantation development (Harrison et al., 2017; Shao et al., 2017a, 2017b; Sozen et al., 2018; Zheng et al., 2019) and gastrulation (Beccari et al., 2018; van den Brink et al., 2014). Although embryonic organoids have been shown to mimic many of the morphological and transcriptional hallmarks of the early embryo, their potential to undergo organogenesis has not yet been explored.

The heart is the first organ to form and function in the embryo. Short after gastrulation, heart progenitors are specified, progressively localize anteriorly and organize in a crescent-shaped domain (around E7.5), the first cardiac compartment that is morphologically identifiable during development. Cardiogenesis is based on the interaction of two different types of progenitors, namely the first heart field (FHF) and the second heart field (SHF) progenitors, the latter originating from pharyngeal mesoderm (Kelly et al., 2014). The cardiac crescent, mainly formed by FHF progenitors, subsequently rearranges to form a linear heart tube (around E8.0-E8.5) which is the first beating structure. Successively, SHF progenitors contribute to heart tube elongation and the heart undergoes looping, ballooning and septation, giving rise to the four-chambered structure typical of adulthood (Harvey, 2002). Heart organogenesis requires cardiac progenitors to interact with surrounding tissues through mechanical interactions and the secretion of cardiac-inducing factors (Miquerol and Kelly, 2013). These involved tissues especially include the endothelium (Brutsaert, 2003; Brutsaert et al., 1998; Narmoneva et al., 2004) and foregut (Hosseini et al., 2017; Kidokoro et al., 2018; Lough and Sugi, 2000; Nascone and Mercola, 1995; Schultheiss et al., 1995; Varner and Taber, 2012).

We hypothesized that due to their embryo-like multi-axial organization and gene expression patterns, mouse gastruloids (Beccari et al., 2018; van den Brink et al., 2014) could offer a suitable template for studying early heart development because they potentially preserve the crucial tissue-tissue interactions required for this organogenesis. Indeed, studying heart organogenesis in such a complex and spatially organized system could allow the modeling of developmental events in an embryo-like context, where cardiac cells are naturally exposed to the influence of other tissues.

Here we show that self-organizing mouse embryonic stem cells (mESCs) can capture early heart organogenesis *in vitro* with a surprising temporal and spatial accuracy. Exposing small ESC aggregates to a cocktail of three cardiogenic factors in gastruloid culture conditions promotes cardiac development *in vitro* starting from *Mesp1*^*+*^ progenitors, which progressively become restricted to the anterior portion of the gastruloid. Through a combination of light-sheet and confocal microscopy, RNAscope imaging, and FACS, we demonstrate that these embryoids support the formation of *Flk1*^*+*^ cardiovascular progenitors, the generation of a vascular-like network, and the formation of progenitors with a first and second heart field identity. Strikingly, we find evidence for the morphogenesis of an anterior cardiac crescent-like domain, which subsequently gives rise to a beating compartment exhibiting Ca^2+^-handling properties compatible with functional fetal cardiomyocytes. Morphogenesis was established in close spatial proximity to the most anterior portion of a co-developing gut–tube-like structure, which was separated from the cardiac domain by an endocardial-like layer. Therefore, this *in vitro* model of cardiac organogenesis uniquely captures interactions between embryonic tissues in the context of a spatially organized embryo-like entity.

## Results

### Optimization of culture conditions to promote efficient beating portions in gastruloids

Gastruloids occasionally formed a beating domain that is exclusively located within their anterior region (38.5 ± 29.3% formed at 168 h) when cultured for 144 h or longer in N2B27 medium (**Fig. 1A**). The location and activity of this beating structure suggested that it might correspond to a cardiac primordium. We tested whether well-known cardiogenic factors (Rajala et al., 2011) could increase the frequency of this event by adding basic fibroblast growth factor, ascorbic acid, and vascular endothelial growth factor 165 (VEGF) (Kattman et al., 2006), singly or in combination, and we increased nutrient and growth factor availability through volume optimization and shaking (**Fig. 1A, Fig. S1A**). In these culture conditions (termed N2B27+++), the frequency of beating gastruloids increased by more than a factor of two (87.2 ± 15.6% formed at 168 h) (**Fig. 1A, B** and **Supplemental Movie 1**). We noticed that exposure to cardiogenic factors was most effective when applied in combination and between 96 and 144 h (**Fig. S1A-F**), so we kept this protocol for the following experiments. Importantly, culturing in N2B27+++ did not alter the polarization of the gastruloids, the extent of their elongation (**Fig. S1I-J**), nor the timing of the emergence of the beating domains (**Fig. 1A, Fig. S1B-D**) compared to standard conditions. Staining for Gata4 and cardiac troponin T (cTnT) confirmed that the beating structure was cardiac-like (**Fig. 1C**).

**Fig. 1.**
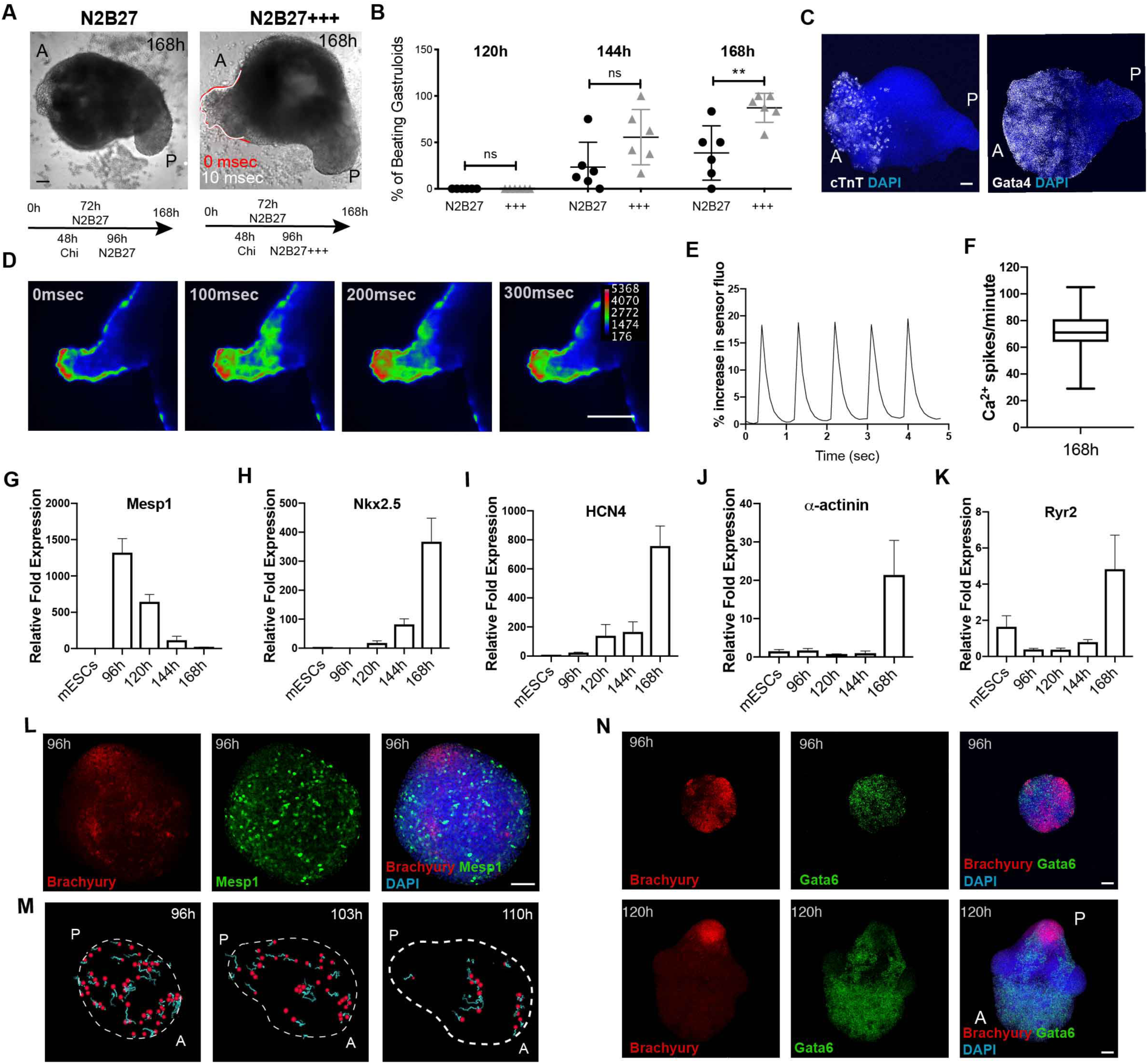
Embryonic organoids recapitulate heart organogenesis. **A**, Gastruloids form a beating portion on their anterior side at 168 h. White and red lines highlight the displacement of the beating domain over 10 msec. **B**, Frequency of beating structures at different time points in *n* = 6 independent experiments. **C**, Immunofluorescence for Gata4 and cTnT on gastruloids at 168 h. **D**, Calcium imaging **E**, representative spiking profile and **F**, spiking frequency of the gastruloid cardiac portion at 168 h for *n* = 21 gastruloids. **G-K**, qPCR gene expression profiles of cardiac genes in gastruloids from 96–168 h. Data are expressed as relative fold expression compared to mESCs in *n* = 4 independent experiments. **L**, Spatial localization of *Mesp1*^*+*^ cells (stained for GFP) at 96 h as compared to Brachyury expression. **M**, Tracking of *Mesp1+* cells from 96–110 h. Red: tracked cells; light blue: cell tracks forward. **N**, Spatial localization of *Gata6*^*+*^ cells at 96 and 120 h compared to Brachyury expression. Scale bars, 100μm. A: anterior; P: posterior. +++: N2B27 with cardiogenic factors.

In cardiomyocytes, physical contraction is coupled to electrical excitation through intracellular changes of Ca^2+^ (Tyser et al., 2016). To evaluate the functionality of the gastruloid cardiac domain, we thus assessed calcium transients, momentary spikes in voltage, by live gastruloid imaging via light-sheet microscopy (at 168 h). Image analysis revealed rhythmic calcium spiking in beating areas with a frequency comparable to beating rates observed in embryos from the crescent stage to the linear heart tube (Tyser et al., 2016) (**Fig. 1D-F, Supplemental Movie 2**). Drugs interfering with calcium transport, including the l-type calcium channel blocker nifidepine and the β-adrenergic agonist isoproterenol, either completely inhibited these transients (**Fig. S2A**) or caused an increase in spiking frequency (**Fig. S2B, C**), respectively. These data demonstrate that the cardiac compartments found in late gastruloids (168 h) exhibit Ca^2+^-handling properties compatible with functional cardiomyocytes.

### The formation of a cardiac domain mimics *in vivo* heart development

To understand if the formation of the cardiac portion mimics developmentally relevant processes, we first analyzed the temporal expression of key genes involved in cardiovascular specification (**Fig. 1G-K**). Similar to what happens during embryonic development (Lescroart et al., 2014; Saga et al., 1996), the first upregulated cardiac gene was *Mesp1*, which was expressed around 96 h and then rapidly downregulated (**Fig. 1G, Fig. S2D, E**). Using a *Mesp1-GFP* reporter ESC line (Bondue et al., 2011) and live light-sheet imaging, we observed that *Mesp1* expression started in a mosaic-like manner at 96 h, with *Mesp1*-positive cells first confined to the anterior side before slowly disappearing after 120 h (**Fig. 1M, Supplemental Movie 3**). At the same time, and concurrently with the elongation and formation of an anterior-posterior axis, *Gata6*-expressing cells localized to the anterior side in gastruloids generated from *Gata6*-*Venus* ESCs (Freyer et al., 2015), opposite to the pole that was positive for Brachyury (**Fig. 1N**). *Gata6* expression was maintained over time (**Fig. S2F, G**). From this stage onwards, the early differentiation genes *Nkx2-5* and *HCN4* were increasingly expressed (**Fig. 1H,I**), followed by genes marking mature cardiomyocytes *(α-actinin, Ryr2*) (**Fig. 1J, K**). This sequence of gene expression shows that gastruloids, stimulated with cardiogenic factors, recapitulate the temporal and spatial gene expression dynamics of cardiac development from the specification of cardiac progenitors to the formation of a beating cardiac structure.

### Co-development of a vascular compartment within the cardiac domain

During embryonic development, the continuous cross-talk of endothelial cells in the developing heart is a prerequisite for cardiomyocyte maturation, function, and survival (Brutsaert, 2003). For this reason, we tested whether such tissue-tissue interactions could potentially take place in developing gastruloids, focusing on cardiovascular progenitors expressing the well-known marker *Flk1* (also known as *Kdr* or *Vegfr2)* (Kattman et al., 2006). In 96-h gastruloids derived from a *Flk1*-*GFP* reporter ESC line (Jakobsson et al., 2010), *Flk1* was expressed at the anterior pole opposite to Brachyury (**Fig. 2A**). Over time, *Flk1* expression persisted in the anterior portion of the gastruloids (**Fig. 2B, Fig. S3A, B**), and *Flk1*-positive cells started to form a vascular-like network of spindle-shaped cells (**Fig. 2C, Supplemental Movie 4**) that stained positive for the endothelial marker CD31 (**Fig. 2D, Fig. S3C**). In an *in vitro* angiogenesis assay, *Flk1*-positive cells that were isolated by fluorescence-activated cell sorting (FACS) from 168-h gastruloids and plated on Matrigel formed vascular-like networks similar to human umbilical vein endothelial cells (HUVECs) (**Fig. 2E, Fig. S3D**). Collectively, these results suggest that gastruloids comprise regions that develop into a vascular-like compartment, which is associated with undergoing cardiac development.

**Fig. 2.**
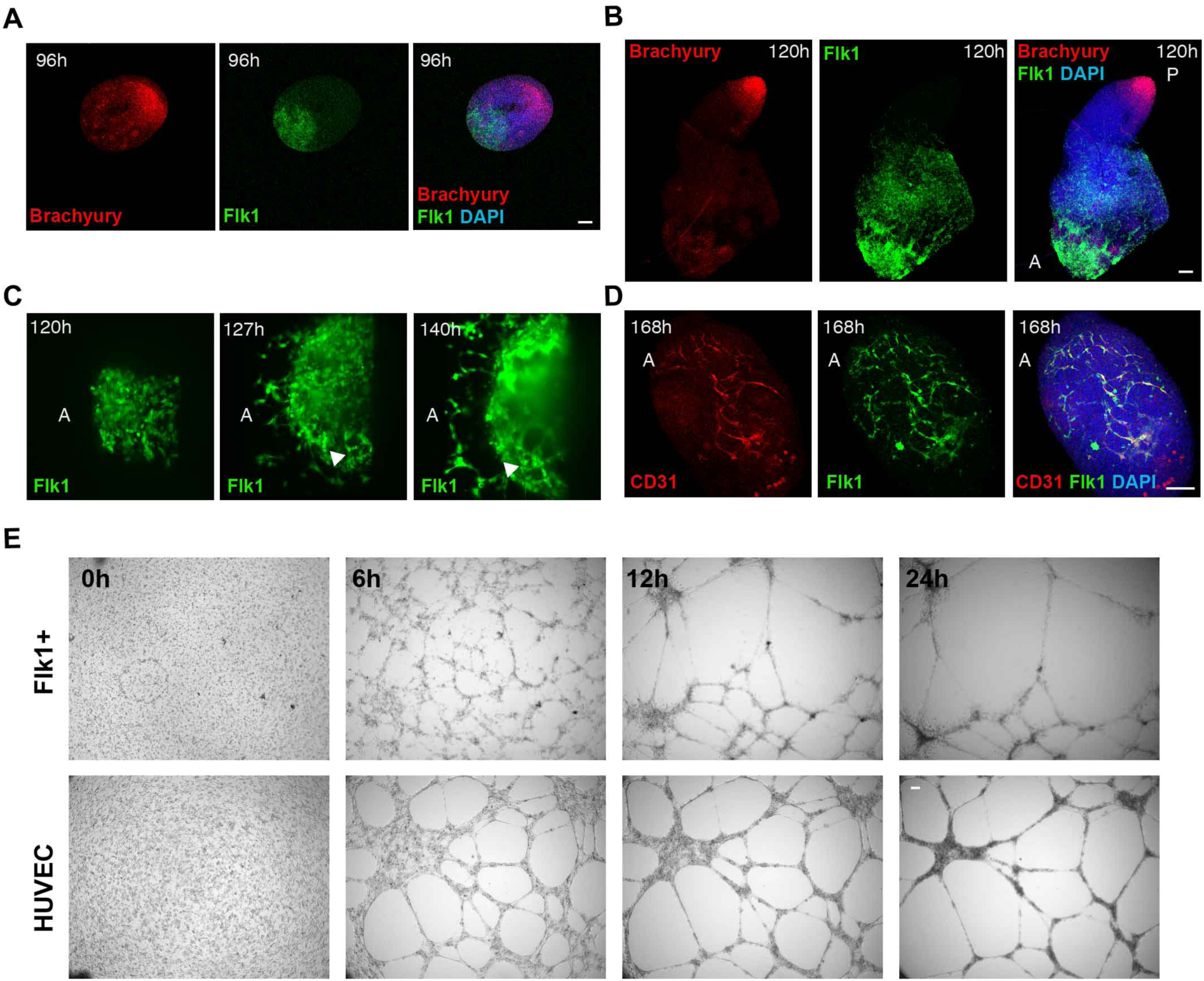
Development of a vascular-like network. Spatial localization of *Flk1*^*+*^ cells at (**A**) 96 and (**B**) 120 h compared to Brachyury expression. **C**, Light-sheet live imaging of the anterior portion of *Flk1-GFP* gastruloids from 120 to 144 h, highlighting the formation of a vascular-like network. **D**, The *Flk1*^*+*^ vascular-like network is positive for CD31. **E**, Angiogenesis assay showing tube formation in isolated *Flk1*^*+*^ cells compared to HUVEC. Scale bars, 100μm. A: anterior; P: posterior.

### *In vitro* cardiac development entails first and second heart field development

A key feature of cardiogenesis is a coordinated interaction between two distinct mesodermal progenitor populations: the FHF, which contributes to the left ventricle and part of the atria, and the SHF, which gives rise to the outflow tract, right ventricle, and part of the atria (Harvey, 2002; Miquerol and Kelly, 2013). After migration from the posterior primitive streak, FHF progenitors form the cardiac crescent and early heart tube, located anteriorly. SHF progenitors, originating from cardiopharyngeal mesoderm (Cortes et al., 2018), are characterized by delayed differentiation and are located medially to the crescent, which can then be involved in heart-tube elongation. Each of these populations is characterized by a specific pattern of gene expression that we also see in our embryonic organoids; we observed the expression of FHF (*Tbx5*) and SHF (*Tbx1*) markers (**Fig. 3A-C**) in mutually exclusive cell populations, and *Isl1* mostly overlapping with *Tbx1* expression (**Fig. 3D**). To corroborate the presence of progenitor populations from both heart fields in our organoids, we used a recently published protocol (Andersen et al., 2018) for FACS-isolating SHF progenitors based on the expression of the C-X-C chemokine receptor type 4 (CXCR4) (**Fig. 3E**). We observed that *Gata6*^*+*^*/*CXCR4^+^ cells express higher levels of SHF markers (*Tbx1, Isl1*, and *FGF10*), but low or unchanged levels of FHF markers (*Nkx2-5, Tbx5*, and *HCN4*) compared to *Gata6*^*+*^*/*CXCR4^-^, confirming their SHF identity (**Fig. 3F-K**). Together, these data show that late gastruloids contain key cell types that are involved *in vivo* in first and second heart–field-based cardiogenesis.

**Fig. 3.**
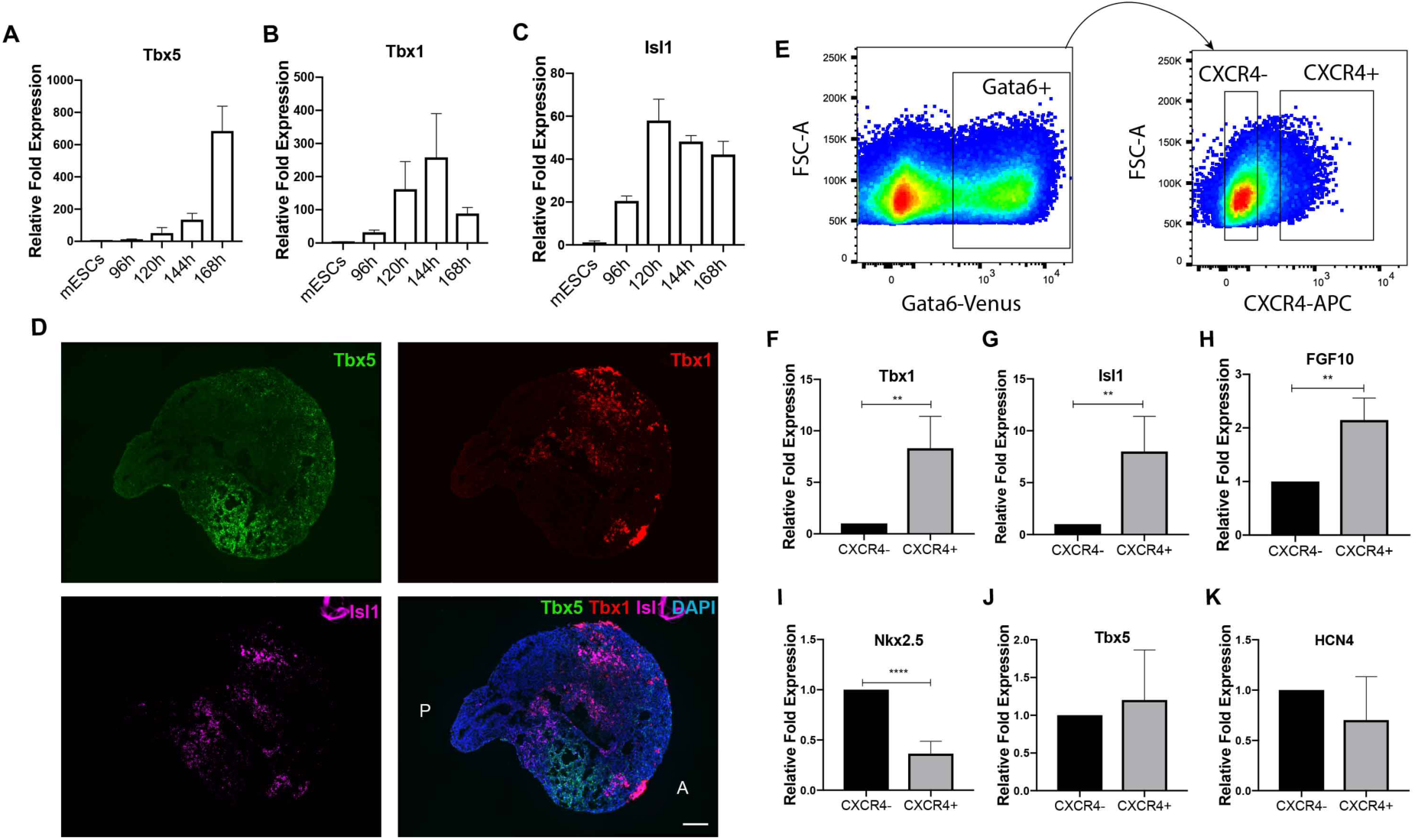
Embryonic organoids form a first and second heart field. Expression levels of markers of (**A**) FHF and (**B, C**) SHF in gastruloids from 96 to 168 h. Data are expressed as relative fold expression compared to mESCs in *n* = 4 independent experiments. **D**, RNA-scope showing spatial localization of FHF and SHF domains. **E**, Representative plot showing the gating strategy used to isolate SHF-enriched progenitors. Expression levels of markers of (**F-H**) SHF and (**I-K**) FHF in *Gata6*^+^/CXCR^-^ and *Gata6*^+^/CXCR^+^ cells isolated from gastruloids at 168 h. Data are expressed as relative fold expression compared to *Gata6*^+^/CXCR^-^ cells in *n* = 4 independent experiments. Scale bar, 100μm. A: anterior; P: posterior.

### *In vitro* cardiac development occurs through establishment of a crescent-like structure

After their specification in the mouse embryo, cardiac progenitors migrate antero-laterally and progressively fuse at the midline to define the first morphologically identifiable heart structure, the cardiac crescent, which appears around E7.5 (Miquerol and Kelly, 2013). Then, morphogenetic movements associated with foregut closure form a linear heart tube by E8.5 (Miquerol and Kelly, 2013). We explored whether gastruloids stimulated with cardiogenic factors could capture these morphological hallmarks of cardiogenesis. Remarkably, in gastruloids cultured for 144 h to 168 h, we observed a recapitulation of these events. Around 144 h, cTnT-positive cardiomyocytes were organized in crescent-like domains (**Fig. 4A**). These further developed into denser crescent-like structures with beating areas, leading to beating epithelial protrusions on the anterior portion of the gastruloids at around 168 h. Similar to mouse embryos (Ivanovitch et al., 2017; Le Garrec et al.; Tyser et al., 2016), these phenotypes could be observed in gastruloids within 24 h, and could be aligned to progressive morphological stages of embryonic cardiac development between E7.5 and E8.5 (**Fig. 4B**). Indeed, a comparative volume analysis of the cTnT-positive gastruloid domains with defined artificial shapes revealed a gradual transition from an almost spherical to a crescent-like shape (144 to 168 h) that then became concave (168 h) (**Fig. 4C**). These results highlight the remarkable capacity of embryonic organoids to promote the spatially and temporally orchestrated morphogenetic processes involved in the early stages of heart development.

**Fig. 4.**
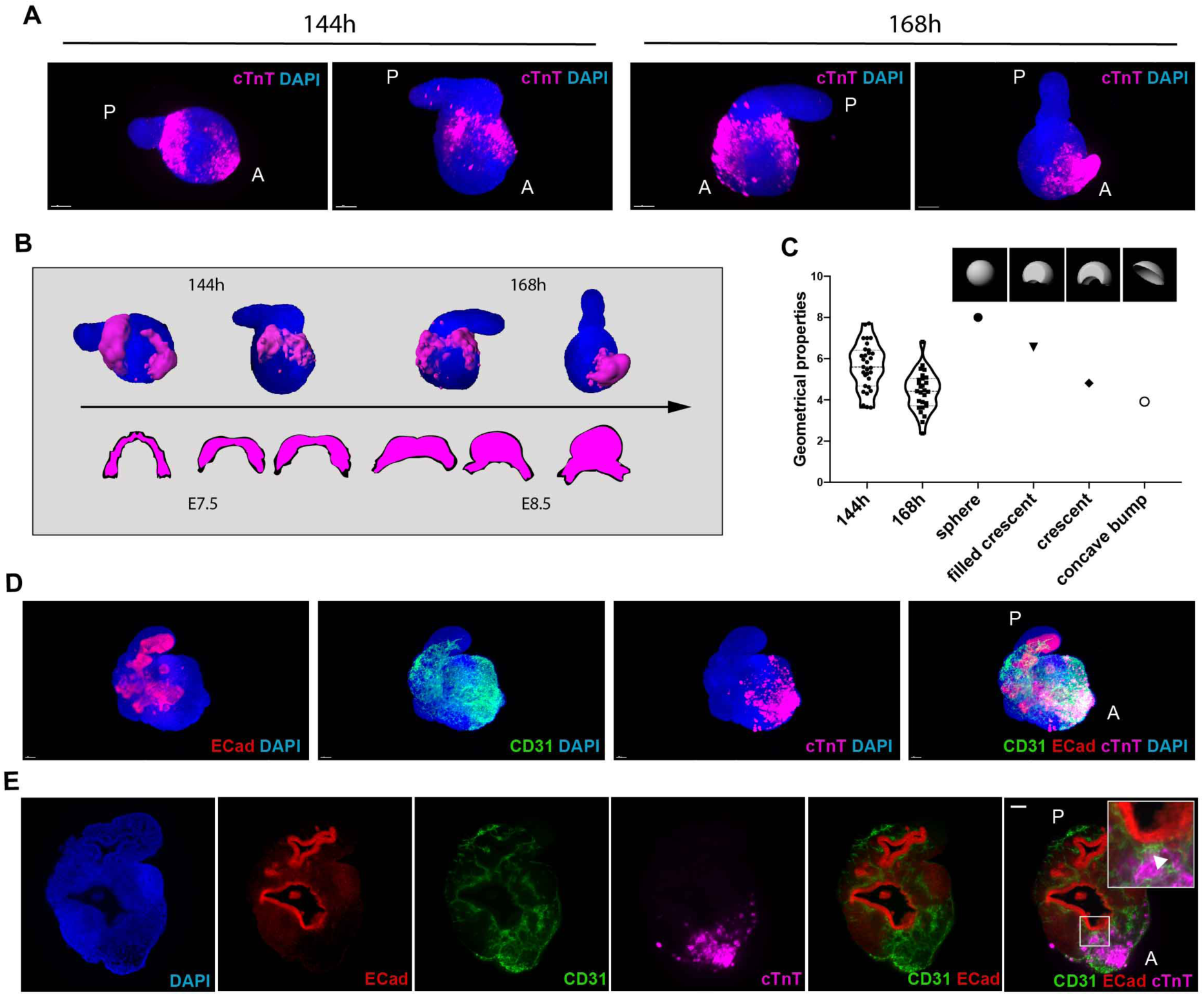
Embryonic organoids recapitulate cardiac morphogenesis. **A**, Light-sheet imaging of cleared gastruloids from 144–168 h show an initial crescent-like domain that is condensed into a beating bud at 168 h. **B**, Schematic illustrating the comparison between gastruloid stages of cardiac development and embryonic stages from cardiac crescent to linear heart tube. **C**, Quantification of geometrical properties (spareness) of the gastruloid cardiac domains compared to those of defined artificial shapes, with *n* = 31 gastruloids at 144 h and *n* = 27 gastruloids at 168 h. **D, E**, The cardiac domain is localized near the anterior epithelial gut–tube-like structure, separated by a CD31^+^ endocardial layer. Scale bar, 100μm. A: anterior; P: posterior.

### *In vitro* cardiogenesis happens in the context of physiological tissue-tissue interactions

The presence of a crescent-like structure and its evolution to a coherent beating group of cells led us to test whether this was accompanied by an association between the crescent and anterior endoderm, as is the case in the embryo (Ivanovitch et al., 2017; Lough and Sugi, 2000). Bulk transcriptomics analysis of late gastruloids revealed gene expression patterns (Sox17, Shh, CDX2) that suggested the presence of a primitive gut-tube-like tissue (Beccari et al., 2018). Strikingly, we found that the cTnT-positive cardiac domain in 168 h gastruloids was exclusively located next to a tube-shaped E-Cadherin-positive epithelial tissue (n = 14/14), with a CD31-positive, putative endocardial-like layer in between (**Fig. 4D,E, Supplemental Movie 5**). This is highly reminiscent of the spatial arrangement of anterior structures in the developing embryo (Ivanovitch et al., 2017).

## Discussion

The heart develops through complex interactions between cardiac progenitors and surrounding tissues (Miquerol and Kelly, 2013). These interactions are crucial for sustaining cardiogenesis but have thus far not been reproduced *in vitro*. Indeed, numerous *in vitro* cardiac models, such as engineered heart muscles (Huebsch et al., 2016; Lind et al., 2017; Ma et al., 2015; Mills et al., 2019; Zhao et al., 2019) or cardiac spheroids (Giacomelli et al., 2017; Polonchuk et al., 2017), have been developed, though the *in vitro* modeling of heart morphogenesis has been out of reach. Our results demonstrate that embryonic organoids can be stimulated to recapitulate *in vitro* the key steps of early cardiac development that *in vivo* require polarized interactions between different primordia. These interactions are critical for achieving the architectural and compositional complexity that are absent in conventional organoids (Rossi et al., 2018), but appear to be provided by the multi-axial properties and spatial organization of the different embryonic tissues that are characteristic of gastruloids (Beccari et al., 2018).

We show that embryonic organoids can be stimulated to form a cardiac portion that involves key developmental hallmarks such as the generation and spatial arrangement of early cardiac progenitors and first and second heart field compartmentalization. Of note, precardiac spheroids were previously shown to generate progenitors with FHF and SHF identity, however, these spheroids did not develop an organized structure (Andersen et al., 2018). In contrast, in our model system, these populations emerged in an embryo-like, spatially organized structure. Moreover, the cardiac portion in our system co-developed with a vascular network and an endodermal component, both known to influence cardiac development (Brutsaert, 2003; Lough and Sugi, 2000). Due to the importance of these neighboring developing tissues, previous attempts were performed to derive spheroids composed of both endothelial and cardiac cells (Giacomelli et al., 2017; Polonchuk et al., 2017). However, these systems, based on an aggregation of cells pre-differentiated in 2D, lacked any visible structural organization. Future work will explore whether the morphogenetic processes seen in our system require the presence of spatial complexity (e.g. multi-axial embryonic patterning) and crosstalk with other tissues.

Collectively, our data show that embryonic organoids have the potential for modeling organogenesis. Herein, we have steered the self-organization of small aggregates of mESCs to form a cardiac primordium as a precursor of an embryonic heart. In the future, it will be interesting to study whether cardiac primordia can be further developed and matured, and what type of tissue-tissue interactions are functionally involved in their development. Additionally, the principles presented here should be applicable to other organ systems once it is possible to overcome the significant challenge of the limited lifespan of embryonic organoids.

## Acknowledgements

We thank Alfonso Martinez Arias for great support throughout the project. We thank Denis Duboule and Nadia Mercader for useful feedback on the manuscript. We thank members of the Lutolf laboratory for discussions and sharing materials, Cédric Blanpain (ULB) for providing *Mesp1-GFP* cells, Alexander Medvinsky (MRC Edinburgh) for providing *Flk1-GFP* cells, Andy Oates (EPFL) and Peter Strnad (Viventis Sàrl) for making light-sheet LS1 microscope available and for technical support, Olivier Burri for EasyXT development, Thierry Laroche for support with Z1 light-sheet imaging, Arne Seitz and other members of Bioimaging and Optics Facility (EPFL) for microscopy support, Jessica Sordet-Dessimoz and Gian-Filippo Mancini from Histology Core Facility for RNAscope, and all personnel of Histology Core Facility, Flow Cytometry Core Facility and Gene Expression Core Facility (EPFL) for their technical support. This work was funded by EPFL.

## Author Contributions

G.R. and M.P.L. conceived the study, designed experiments, analyzed data and wrote the manuscript. G.R. performed the experiments, A.B. performed *in vivo* light-sheet imaging, R.G. developed the script for the analysis of cardiac structures, M.G. contributed to protocol optimization and data analysis, R.G.K. contributed to study design and data discussion and provided feedback on the manuscript.

## Materials and Methods

### Cell Culture

mESCs were cultured at 37°C, 5%CO_2_ in DMEM supplemented with 10% Embryonic Stem Cell qualified FBS (Gibco), NEAA, Sodium Pyruvate, β-mercaptoethanol, 3µM CHI99201 (Chi), 1µM PD025901 and 0.1µg ml^-1^ LIF. *Gata6-Venus* (Freyer et al., 2015), *Flk1-GFP* (Jakobsson et al., 2010), and *Mesp1-GFP* (Bondue et al., 2011) cells were cultured on gelatin-coated tissue culture flasks; *Sox1-GFP::Brachyury-mCherry* (Deluz et al., 2016) cells on tissue-culture flasks without coating. If not differently specified, *Sox1-GFP::Brachyury-mCherry* cells were used for our experiments. HUVECs were cultured in EGM-2 medium (Lonza). All cells were routinely tested for Mycoplasma with Mycoalert mycoplasma detection kit (Lonza) or by PCR.

### Gastruloid culture

Gastruloids were generated as previously described (Baillie-Johnson et al., 2015). Briefly, 300-700 mESCs were plated in 40μl N2B27 in 96-well Clear Round Bottom Ultra-Low Attachment Microplates (7007, Corning). After 48 h, 150μl of N2B27 containing 3µM Chi were added to each well. After 72 h, medium was changed with N2B27. Starting from 96 h, the protocol was optimized as described in **Fig. S1A**. At 96 h, gastruloids were transferred in Ultra-Low Attachment 24-well Plates (3473, Corning) in 100μl of medium, plus 700μl of fresh N2B27 containing 30ng ml^-1^ bFGF (PMG0034, Gibco), 5ng ml^-1^ VEGF 165 (PHC9394, Gibco) and 0.5mM L-ascorbic acid phosphate (013-12061, Wako) (N2B27+++) and cultured on an orbital shaker placed at 37°C, 5%CO_2_ at 100rpm (VWR mini shaker). From 120 h onwards, half medium was changed daily. Unless differently specified, N2B27+++ was applied from 96 to 144 h, while from 144 h to 168 h N2B27 was used for medium change.

### Live imaging and cell tracking

Bright-field live imaging of beating gastruloids was performed with a Nikon Ti inverted microscope equipped with an incubation chamber at 37°C, 5%CO2. Light-sheet live imaging of *Flk1-GFP* and *Mesp1-GFP* gastruloids was performed with a prototype of LS1 live inverted light-sheet microscope (Viventis Microscopy Sarl, Switzerland), at 37°C, 5% CO_2_. A volume of 150-200µm was acquired with a Z spacing of 2-3µm between slices and pictures were captured every 20 min for *Flk1-GFP* gastruloids and every 10 min for tracking of *Mesp1*^+^ cells. *Flk1-GFP* light-sheet video montages were obtained with the Arivis Vision4D software. To track *Mesp1*^*+*^ cells in Gastruloids from 96 to 120 h, LS1 live light-sheet images were processed with the Fiji Mastodon plugin, using a semi-automatic tracking. Subsequently, Mastodon files were exported for Mamut, and the Fiji Mamut plugin was used to display cell tracks as shown in **Figure 1**.

### Immunofluorescence, confocal and light-sheet imaging on fixed samples

Immunofluorescence on whole mount gastruloids was performed as previously described (Baillie-Johnson et al., 2015). Briefly, gastruloids were washed in PBS and fixed in 4% PFA for 2 h at 4°C while shaking. Samples were washed 3 times in PBS and 3 times (10 min each) in blocking buffer (PBS, 10%FBS, 0,2%Triton X-100), then blocked for 1 h at 4°C in blocking buffer. Gastruloids were then incubated O/N with primary antibodies in blocking buffer, at 4°C while shaking. The day after, gastruloids were washed 4 times (20 min each) with blocking buffer, at 4°C while shaking, and incubated O/N with secondary antibodies and DAPI (2µg ml^-^ 1, Sigma-Aldrich) in blocking buffer, at 4°C while shaking. The day after, gastruloids were washed for 1h with blocking buffer, at 4°C while shaking, then rinsed in PBS, 0,2%FBS, 0,2% Triton X-100 and mounted on Superfrost plus glass slides (ThermoFisher) with Floromount-G for confocal imaging (Southern Biotech). The following primary antibodies were used: mouse anti-Gata4 (1:500, Santa Cruz Biotechnology, G-4); chicken anti-GFP (1:750, Aves Labs); goat anti-Brachyury (1:300, Santa Cruz Biotechnology, C-19); rat anti-CD31 (1:100, BD, MEC 13.3), mouse anti-cardiac troponin T (1:100, ThermoFisher, 13-11), rabbit anti-E-Cadherin (1:500, Cell Signaling, 24E10). The following secondary antibodies were used: donkey anti-chicken 488 AlexaFluor (1:500, Jackson ImmunoResearch); donkey anti-goat AlexaFluor 568 (1:500, ThermoFisher); goat anti-rat AlexaFluor 568 (1:500, ThermoFisher); goat anti-mouse AlexaFluor 647 (1:500, ThermoFisher); donkey anti-rabbit 568 (1:500, ThermoFisher). Confocal pictures were acquired with a Zeiss LSM 700 inverted confocal microscope equipped with a Axiocam MRm black and white camera in the EPFL bioimaging and optics facility. For light-sheet imaging (**Fig. 4**), samples were mounted in 1% low-melt agarose and cleared overnight with CUBIC mount solution (Lee et al., 2016). Light-sheet imaging was performed on a Zeiss Light-sheet Z1 microscope equipped with a Plan-Neofluar 20x/1.0 Corr nd=1.45 objective. Light-sheet images were further processed with Imaris software, for 3D rendering and surface generation.

### RNA extraction and qRT-PCR

RNA was extracted from gastruloids with the RNeasy Micro kit (Qiagen), according to manufacturer’s instructions and quantified with a spectrophotometer (ND-1000, Nanodrop). 1μg of RNA was reverse-transcribed with the iScript cDNA Supermix kit (Biorad). cDNA was diluted 1:10 and 1.5μl of cDNA per reaction were used, in a total volume of 10μl. 384 well plates were prepared using a robotized liquid handling platform (Hamilton Microlab Star). qPCR was run with a 7900HT Fast PCR machine (Applied Biosystems), using Power SYBR Green PCR Master Mix (Applied Biosystems), with an annealing temperature of 60°C. Gene expression was normalized on *β-actin* expression. Relative fold expression was calculated with the 2−ΔΔCT method. 500nM of the following primers were used: *Mesp1* FOR GTCTGCAGCGGGGTGTCGTG; *Mesp1* REV CGGCGGCGTCCAGGTTTCTA; *Nkx2.5* FOR CACATTTTACCCGGGAGCCT; *Nkx2.5* REV ACCAGATCTTGACCTGCGTG; *HCN4* FOR GTGGGGGCCACCTGCTAT; *HCN4* REV GTCGGGTGTCAGGCGGGA; *α-actinin* FOR GGGCTATGAGGAGTGGCTATT; *α-actinin* REV AGTCCTTCTGCAGCAAGATCT; *RyR2* FOR TGCATGAGAGCATCAAACGC; *RyR2* REV CGCGGAGAGAGGCATTACAT; *Tbx5* FOR GGCATGGAAGGAATCAAGGTG; *Tbx5* REV TTTGGGATTAAGGCCAGTCAC; *Tbx1* FOR CTGTGGGACGAGTTCAATCAG; *Tbx1* REV TTGTCATCTACGGGCACAAAG; *Isl1* FOR ATGATGGTGGTTTACAGGCTAAC; *Isl1* REV TCGATGCTACTTCACTGCCAG; *FGF10* FOR TCAGCGGGACCAAGAATGAAG; *FGF10* REV CGGCAACAACTCCGATTTCC; *β-actin* FOR CTGTCGAGTCGCGTCCACC; *β-actin* REV CGCAGCGATATCGTCATCCA.

### RNAscope

Gastruloids were washed in PBS and fixed O/N in 4% PFA, at 4°C while shaking. The day after, samples were washed 3 times in PBS and included in HistoGel (ThermoFisher) blocks. HistoGel blocks were then processed with a Tissue-Tek VIP 6 AI Vacuum Infiltration Processor (Sakura) and included in paraffin. Paraffin blocks were cut at 4μm with a Hyrax M25 microtome (Zeiss). RNA-scope was performed with the ACDBio Manual assay kit using RNAscope Probe-Mm-Tbx1 (481911), RNAscope Probe-Mm-Isl1-C3 (451931-C3) and RNAscope Probe-Mm-Tbx5-C2 (519581-C2) probes, according to manufacturer’s instructions. Polr2a-C1, Ppib-C2 and Ubiquitin-C3 probes were used as positive and negative controls. Pictures were acquired with an upright Leica DM5500 microscope equipped with a CCD DFC 3000 black and white camera.

### Flow cytometry analysis and FACS

Gastruloids were collected, washed in PBS, and digested in 4mg ml^-1^ dispase I (Roche), 3mg ml^-1^ collagenase IV (Gibco) and 100μg ml^-1^ DNase I (Roche) in PBS (2 digestion cycles at 37°C, 5 min each; gentle pipetting was applied between the two cycles to mechanically dissociate the gastruloids). Digestion was blocked with DMEM containing 10% FBS, then samples were centrifuged and the cell pellet was resuspended in sorting buffer (PBS, 5%FBS, 1mM EDTA, 1%P/S) for antibody staining. Samples were incubated for 1 h on ice with antibodies, and 30min on ice with Aqua live/dead fixable dead cell stain kit (405/525nm, Invitrogen) or 10 min on ice with DAPI. Unstained, FMO and single color samples were used as controls. The following antibodies were used: anti CD31-PE 1:1200 (BD, MEC 13.3), anti VEGFR2/Flk1-APC 1:200 (Biolegend, Avas12); anti CXCR4-APC 1:100 (BD, 2B11). Samples were analyzed with a BD LSR II flow cytometer. Cell sorting was performed using a BD FACSAria Fusion cell sorter.

### *In vitro* angiogenesis assay

168-h *Flk1-GFP* gastruloids were collected and digested as described above for FACS analysis. *Flk1*^+^ and *Flk1*^*-*^ cells were isolated through cell sorting using a BD FACSAria Fusion cell sorter. 5*10^4^ cells per condition were plated in IBIDI μ-angiogenesis slides pre-coated with 10μl reduced growth factor Matrigel (Corning) in the lower chamber. Undifferentiated mESCs and HUVEC were used as negative and positive controls, respectively. Live imaging of tube formation was performed with a Nikon Ti inverted microscope equipped with an incubation chamber at 37°C, 5% CO_2,_ with acquisitions every 15 min.

### Calcium imaging

To image calcium fluxes, gastruloids were incubated for 1 h with 8μM Cal-520 (AAT Bioquest) at 37°C, 5% CO_2_. Gastruloids were then transferred to fresh medium before imaging. Imaging was performed with a Light-sheet Z1 microscope (Zeiss) equipped with an environmental chamber to maintain gastruloids at 37°C and 5% CO_2_. For imaging, gastruloids were embedded in 1% low melt agarose and the chamber was filled with culture medium. Nifidepine (Sigma Aldrich, 10μM) and isoproterenol (isoprenaline hydrochloride, Sigma I5627, 1μM) were added with a syringe directly to the imaging chamber during acquisition. The analysis of calcium spikes was performed with the Fiji Stacks-plot Z-Axis profile plugin. The baseline intensity was normalized to the minimum value over 10 sec. The ratio of fluorescence intensity to baseline intensity was calculated and results are shown as the percentage of increase over the baseline, which shows the relative changes in intracellular Ca^2+^.

### Crescent analysis

Analysis of crescent geometrical properties was performed using a custom-made Matlab script for Imaris files with a Imaris XT feature and EasyXT. Initially, we generated artificial shapes to be used as reference for defined geometrical metrics. Using a custom script (*crescent_generator.m*), 3D objects were generated to mimic crescent structures with different characteristics. To do so, the script creates two spot objects, makes channels from these spots and finally subtracts one channel to the other. To analyze surfaces, 3D images from non-beating organoids at 144 h and beating organoids at 168 h were acquired with a Light-sheet Z1 microscope (Zeiss) after clearing, as described above. For each individual stack, a surface was created using a dedicated user interface (*GUI_DetectAndAnalyze.m*), defining the object that should be created for each channel (*settings_.m*). Due to variability in background and signal intensity, the threshold (absolute intensity) was adjusted manually for each surface. Using a custom script (*makeCroissantMeasure_final.m*) geometrical measurements were computed and the results exported in csv table. In the graph, we plot the measure of Spareness, which is described as the ratio between the volume of the object and the volume of the best fitted ellipsoid. All scripts and settings used for analysis are available at go.epfl.ch/Crescent_Analysis.

### Statistics

All data shown in column graphs are expressed as mean ± SD, apart from the graph showing Calcium spikes frequency, which is expressed as mean ± whiskers from min to max. All other graphs show single data points. Statistical analysis between two columns was performed using two-tailed unpaired Student’s t test, whereas data containing more than two experimental groups were analyzed with one-way analysis of variance followed by Bonferroni’s test. To calculate the significance of the percentage of increase over baseline frequency after Isoproterenol administration, we applied a one sample t test. Statistical significance was calculated using the Graphpad Prism software, that was also used to generate all graphs. *P < 0.05; **P < 0.01; ***P < 0.001; confidence intervals 95%; alpha level 0.05.

## Supplemental Figures

**Figure S1.**
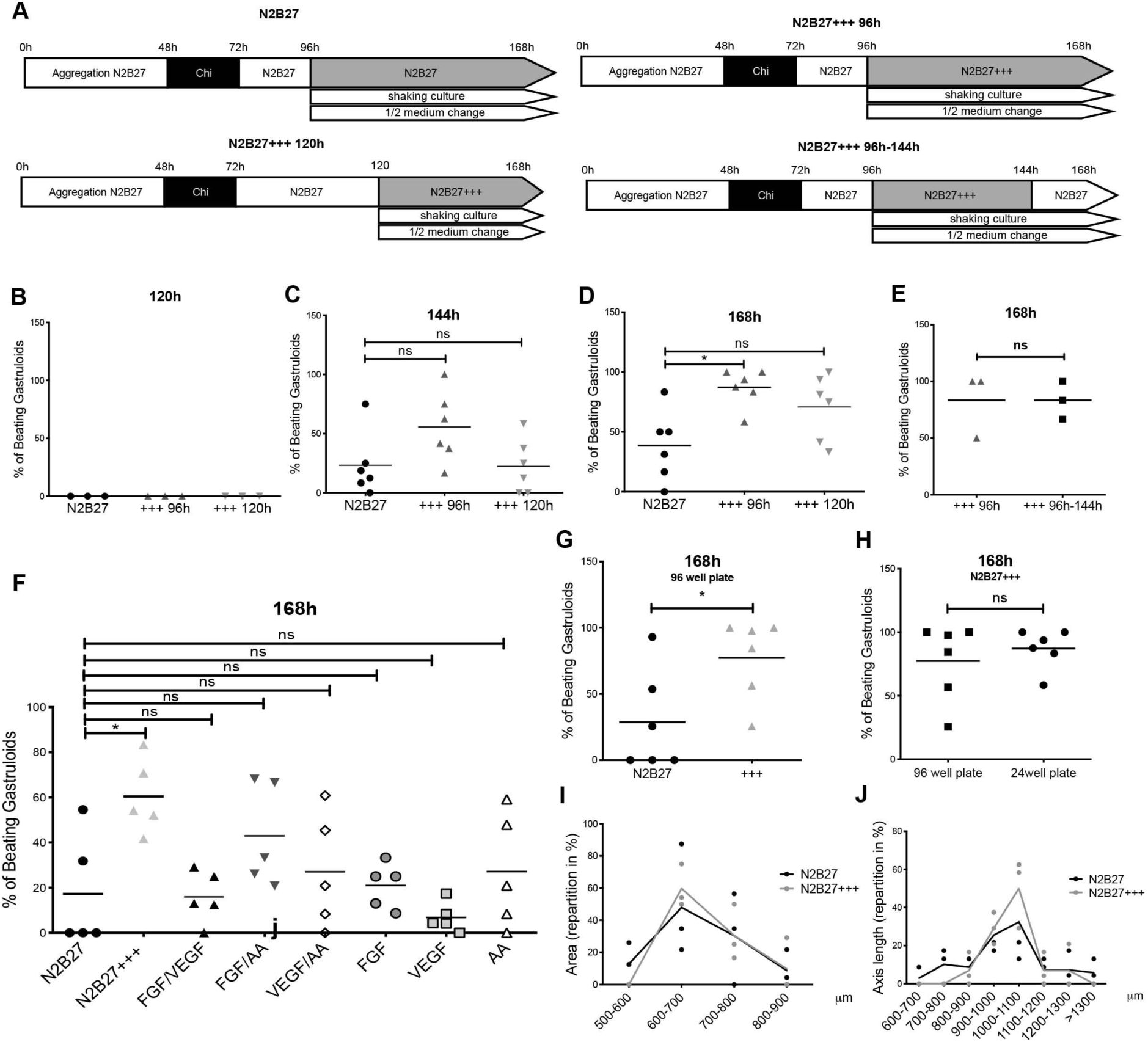
Comparison between N2B27 and N2B27+++ culture conditions. **A**, Schematics of the different culture conditions tested. **B-E**, Percentage of beating gastruloids from 120 to 168h in the different conditions. **F**, Exposure to N2B27+++ induces beating at higher frequencies compared to exposure to single factors or couples. **G, H** Frequencies of beating gastruloids grown in 96well plates (**G**) and comparison with those grown in 24 well plates from 144h (**H**). Each dot in **B-H** represent an independent experiment. **I-J**, quantification of area and axis at 168h of gastruloids grown in N2B27 or N2B27+++. Mean of *n*=3 independent experiments.

**Figure S2.**
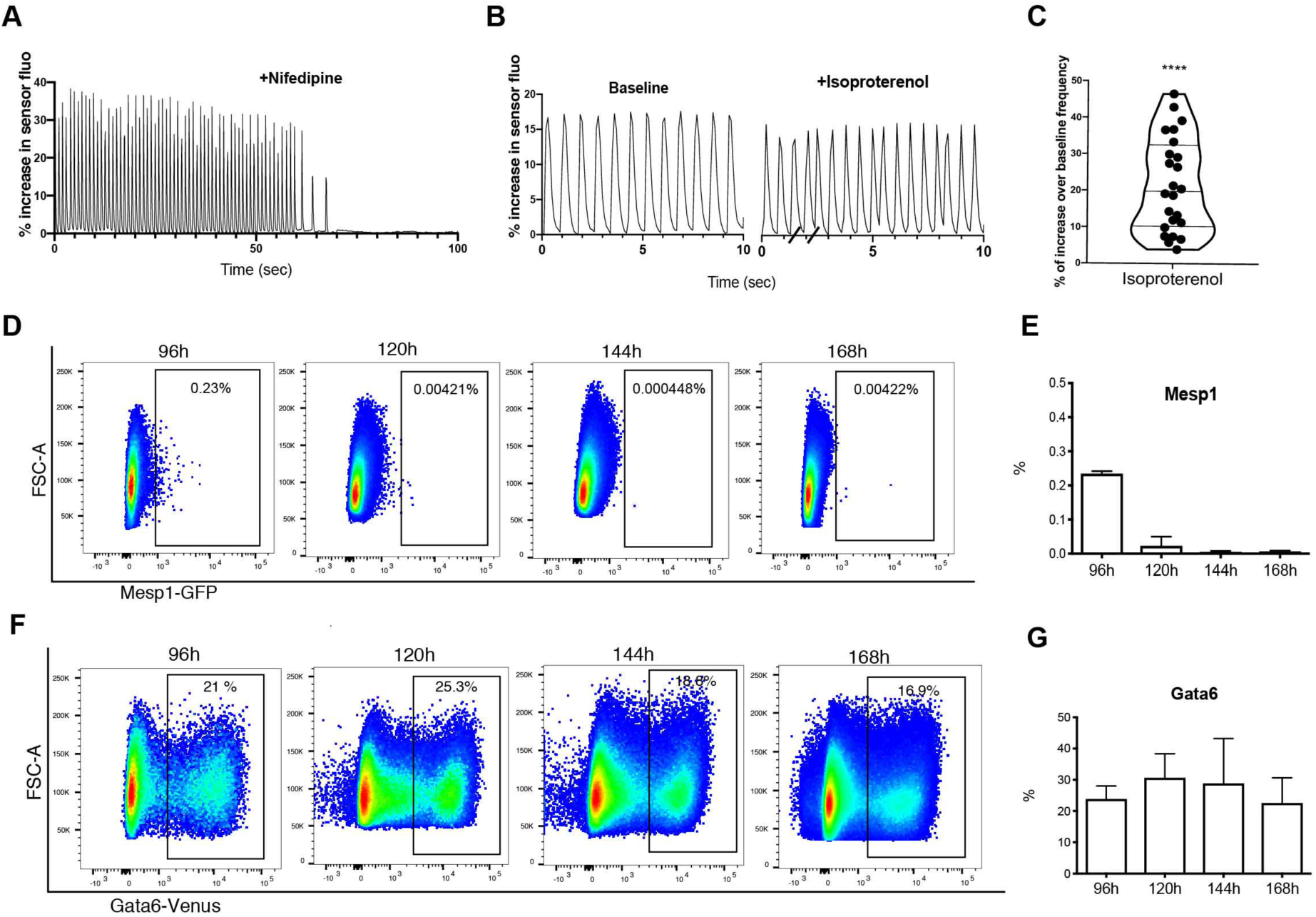
Development of the cardiac portion of gastruloids. **A-C**, Treatment of 168 h gastruloids with Nifedipine (*n*=9 gastruloids) (**A**) or Isoproterenol (**B, C**) abolish or fastens calcium spiking in the gastruloid cardiac portion, respectively. Graph in (**B**) shows representative calcium spikes of gastruloids before and after Isoproterenol treatment, and graph in (**C**) the percentage of increase over baseline frequency of gastruloids after Isoproterenol treatment. *n*=24 gastruloids. **D, E**, FACS analysis of *Mesp1-GFP* gastruloids from 96 to 168 (**D**) and relative quantification (**E**). *n*=2 independent experiments. **F, G**, FACS analysis of *Gata6-Venus* gastruloids from 96 to 168 (**F**) and relative quantification (**G**). *n*=2 independent experiments.

**Figure S3.**
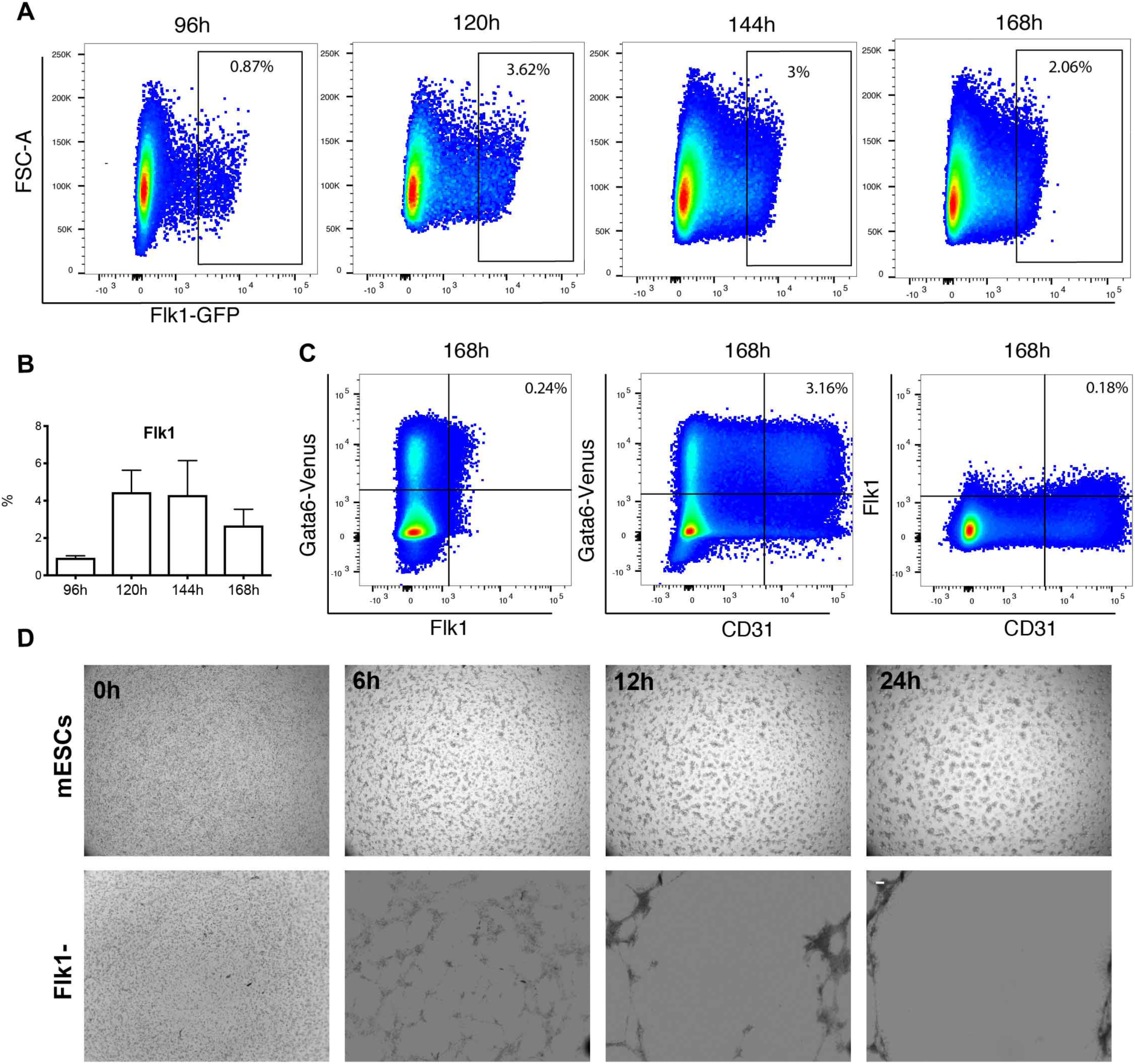
Flk1 marks a vascular-like compartment. **A, B** FACS analysis of *Flk1-GFP* gastruloids from 96 to 168 (**A**) and relative quantification (**B**). *n*=2 independent experiments. **C**, FACS analysis of *Gata6-Venus* gastruloids showing co-expression of Flk1 and CD31. *n*=2 independent experiments. **D**, angiogenesis assay showing inability to form vascular-like tubes by undifferentiated mESCs and *Flk1*^*-*^ cells. Scale bar, 100μm.

## Supplemental Movie Legends

**Supplemental Movie 1 | Example of a non-beating gastruloid cultured in N2B27 and a beating gastruloid cultured in N2B27+++**

**Supplemental Movie 2 | Calcium imaging of a gastruloid beating portion with Cal520**

**Supplemental Movie 3 | *Mesp1***^***+***^ **light-sheet cell tracking and 3D rendering of *Mesp1***^***+***^ **cell tracking from 96 to 120h**

**Supplemental Movie 4 | Development of a *Flk1***^**+**^ **network in embryonic organoids.** Rendering of Light-sheet LS1 live imaging of the anterior portion of a *Flk1-GFP* gastruloid from 120 to 144 h. The video was created and edited with the Arivis Vision4D software.

**Supplemental Movie 5 | Localization of the gut tube, endocardium and cardiac portion in embryonic organoids.** Rendering of the immunostaining in **Figure 4 D, E** showing respective localization of the gut-like tube, the cardiac portion and the endocardial-like layer in 3D. Imaging was performed using Light-sheet Z1 microscope on optically cleared samples. The video was created and edited with Imaris software.

